# Inhibition Of One-Carbon Metabolism In Ewing Sarcoma Results In Profound And Prolonged Growth Suppression Associated With Purine Depletion

**DOI:** 10.1101/2025.04.13.647987

**Authors:** Sara Zirpoli, Noah Copperman, Shrey Patel, Alexander Forrest, Zhanjun Hou, Larry H. Matherly, David M. Loeb, Antonio Di Cristofano

## Abstract

Ewing sarcoma (EWS) is the second most common primary bone malignancy in adolescents and young adults. Patients who present with localized disease have experienced a steadily improving survival rate over the years, whereas those who present with metastatic disease have the same dismal prognosis as 30 years ago, with long term survival rates less than 20%, despite maximal intensification of chemotherapy. Thus, novel treatment approaches are a significant unmet clinical need. Targeting metabolic differences between EWS and normal cells offers a promising approach to improve outcomes for these patients. One-carbon metabolism utilizes serine and folate to generate glycine and tetrahydrofolate (THF)-bound one-carbon units required for *de novo* nucleotide biosynthesis. Elevated expression of several one-carbon metabolism genes is significantly associated with reduced survival in EWS patients. We show that both genetic and pharmacological inhibition of a key enzyme of the mitochondrial arm of the one-carbon metabolic pathway, serine hydroxymethyltransferase 2 (SHMT2), leads to substantial inhibition of EWS cell proliferation and colony-forming ability, and that this effect is primarily caused by depletion of glycine and one-carbon units required for synthesis of purine nucleotides. Inhibition of one-carbon metabolism at a different node, using the clinically relevant dihydrofolate reductase inhibitor Pralatrexate, similarly yields a profound growth inhibition, with depletion of thymidylate and purine nucleotides. Genetic depletion of *SHMT2* dramatically impairs tumor growth in a xenograft model of EWS. Together, these data establish the upregulation of the one-carbon metabolism as a novel and targetable vulnerability of EWS cells, which can be exploited for therapy.

**Statement of Significance:** Using both genetic and pharmacologic approaches, this study identifies Ewing sarcoma’s dependence on the mitochondrial arm, but not the cytoplasmic arm, of one-carbon metabolism as a targetable vulnerability that can be effectively harnessed for therapy.

## Introduction

Ewing sarcoma (EWS) is the second most common bone tumor in children, adolescents, and young adults. Patients who present with localized disease have experienced steady improvements in survival in a series of clinical trials have demonstrated the efficacy of increased chemotherapy intensity (1). In contrast, patients who either present with metastatic disease or who suffer a relapse have the same grim prognosis as in the 1980’s, with no improvement in survival despite numerous clinical trials aimed at maximizing chemotherapy intensity (2). For these patients, it is clear that future improvements in patient outcomes will not derive from modifying their chemotherapy but rather from identifying alternative therapeutic approaches based on a deeper understanding of EWS biology. One promising approach is to identify metabolic differences between normal cells and EWS cells that can be exploited for therapeutic gain.

The primary oncogenic mutation driving EWS is a chromosomal translocation, most frequently t(11;22), resulting in the production of the EWS-FLI1 oncoprotein, that has both transcriptional regulatory and mRNA splicing activity (3). The literature suggests that the EWS-FLI1 fusion oncogene directly regulates the expression of several enzymes involved in de novo serine and glycine biosynthesis and in one-carbon metabolism (4,5). Furthermore, inhibition of serine synthesis at phosphoglycerate dehydrogenase in EWS cells significantly inhibits cell proliferation in the absence of exogenous serine and glycine, in line with a critical role of one-carbon metabolism in supporting EWS cell growth (6). The one-carbon metabolic pathway utilizes serine and dietary folate to generate glycine and tetrahydrofolate (THF)-bound one-carbon units, which are essential for a variety of cellular processes, including de novo nucleotide biosynthesis, NADPH and glutathione production, methionine synthesis, biological methylation and mitochondrial protein translation (7-9). One-carbon metabolism has two compartmentalized arms, one oxidative in the mitochondria, driven by serine hydroxymethyltransferase 2 (SHMT2), 5,10-methylene THF dehydrogenase 2 (MTHFD)2, and MTHFD1-like (MTHFD1L), and one reductive in the cytoplasm, driven by SHMT1 and MTHFD1. These two arms form a normally unidirectional cycle between the two cellular compartments (10) (Fig.1A).

**Figure 1.**
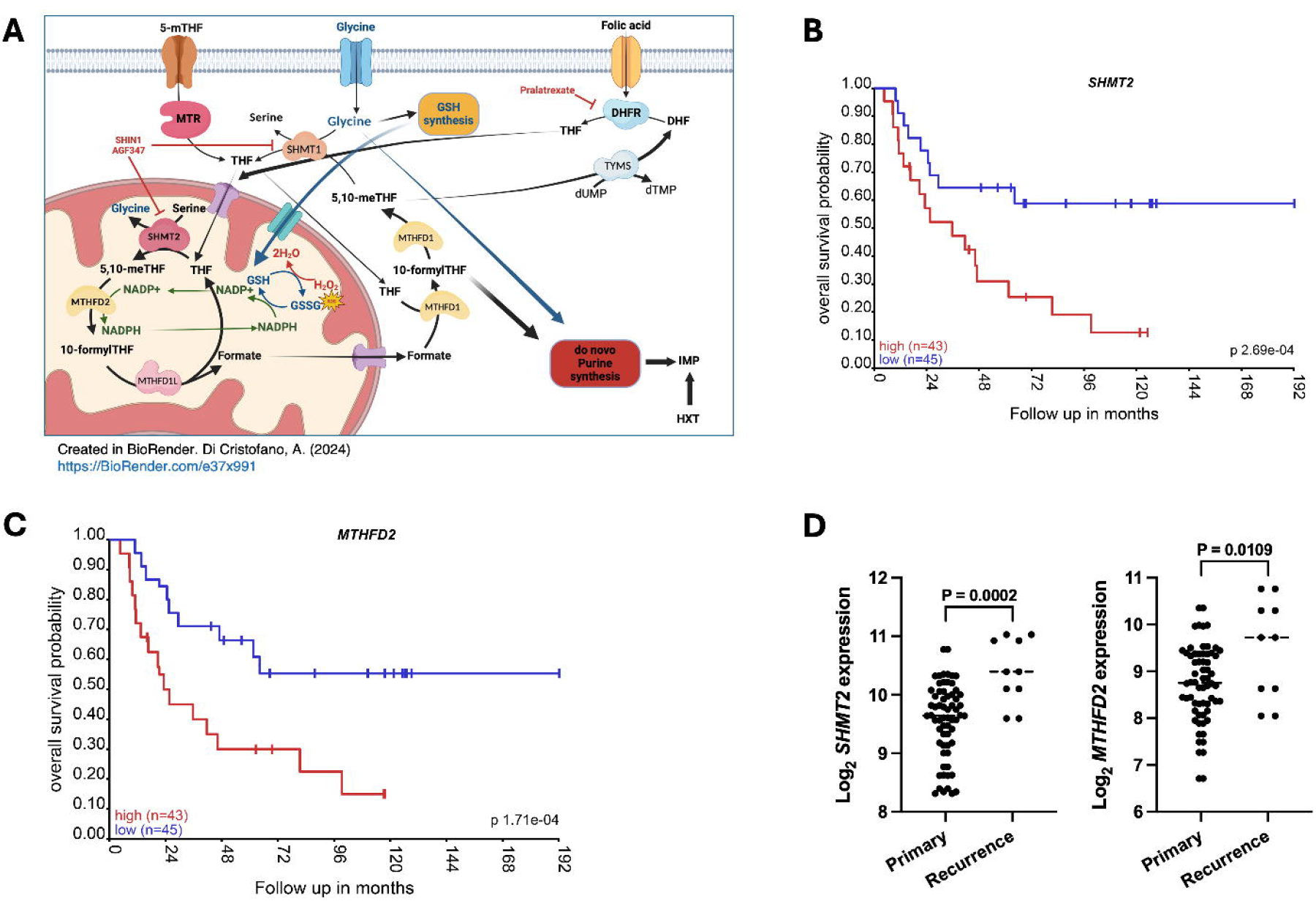
High expression of *SHMT2* and *MTHFD2* is associated with more aggressive clinical features. A) Cartoon depicting the key steps of the cytoplasmic and mitochondrial arms of one-carbon metabolism, as well as the targets of the three inhibitors utilized in this study. MTR: Methionine synthase, DHFR: Dihydrofolate reductase, TYMS: Thymidylate synthase, SHMT: Serine hydroxymethyltransferase, MTHFD: Methylenetetrahydrofolate dehydrogenase, THF: tetrahydrofolate, DHF: Dihydrofolate, 5-mTHF: 5-methyl-tetrahydrofolate, 5,10-meTHF: 5,10-Methylene-tetrahydrofolate, GSH: Glutathione, GSSG: Glutathione disulfide, NADP: Nicotinamide adenine dinucleotide phosphate, IMP: Inosine monophosphate, HXT: Hypoxanthine. B, C) Overall survival of EWS patients (n=88) expressing high or low levels of *SHMT2* (B) or *MTHFD2* (C). D) Expression levels of *SHMT2* or *MTHFD2* in patients with primary (n=64) and recurrent (n=10) disease. Analysis was performed on dataset GSE17679 using R2: Genomics Analysis and Visualization Platform (http://r2.amc.nl).

SHMT2 and MTHFD2 are among the most consistently upregulated metabolic genes in cancer (11,12), supporting the critical role of one-carbon metabolism in the production of metabolites to fuel rapid growth and cell division and to survive intracellular and extracellular oxidative stress. The present study combines genetic and pharmacological approaches to explore the role of one-carbon metabolism in EWS and demonstrates that EWS cells have an absolute dependence on mitochondrial one-carbon metabolism, which is essential for de novo nucleotide biosynthesis and glycine production. Furthermore, we provide compelling in vivo evidence that targeting one-carbon metabolism effectively impairs tumor growth in a xenograft model of EWS.

## Materials and Methods

### Cell Culture

The human EWS cell lines used in this study (Table 1) were maintained at 37°C with 5% CO2 in RPMI1640 (Cytiva) with 10% fetal bovine serum (FBS, Biowest). For all the experiments, cells were grown in RPMI1640 with 10% dialyzed FBS (Biowest). Cell line identity was validated by short-tandem repeat (STR) profiling, and cell lines were screened regularly for mycoplasma contamination.

**Table 1.** Driver mutations in the EWS cell lines used in this study.

|  | **FUSION** | **TP53** | **STAG2** | **CDKN2A** |
| --- | --- | --- | --- | --- |
| **CHLA10 (RRID:CVCL_6583)** | EWS-FLI1 | mut | wt | wt |
| **MHH-ES1 (RRID:CVCL_1411)** | EWS-FLI1 | mut | mut | wt |
| **SK-ES1 (RRID:CVCL_0627)** | EWS-FLI1 | mut | mut | wt |
| **TC32 (RRID:CVCL_7151)** | EWS-FLI1 | wt | mut | mut |
| **TC71 (RRID:CVCL_2213)** | EWS-FLI1 | mut | wt | mut |
| **RD-ES (RRID:CVCL_2169)** | EWS-FLI1 | mut | wt | wt |

### shRNA-mediated gene knock-down

Cells were transduced with lentiviruses encoding shRNAs against *SHMT1* (RHS4430-101028254, RHS4430-101032591, and RHS3979-9602175), and *SHMT2* (RHS3979-9602214 and RHS3979-9602215).

ShRNAs for *SHMT2* were cloned into the Tet-pLKO-puro (Addgene #21915; RRID:Addgene_21915). Lentiviral constructs were packaged in HEK293 cells (RRID:CVCL_0045) transfected using Lipofectamine 2000 (Invitrogen). Viral supernatants were collected 48-72 hours after transfection, combined, and filtered using 0.45 μm Nalgene SFCA syringe filters (Thermo Scientific). Target cells were plated 24 hours before infection and then exposed to viral supernatants supplemented with 8 μg/ml polybrene (Santa Cruz). Infected cell lines were then selected for at least 72 hours with 2 μg/ml puromycin (Corning).

Transduced cell lines were pretreated with 0.25 µg/ml doxycycline hyclate (Sigma Aldrich) for 48 hours prior to plating for proliferation experiments. In all experiments performed on transduced cell lines, doxycycline hyclate was replaced every 48 hrs.

### Drugs and chemicals

The following chemical compounds were used: SHIN1 (Aobius), AGF347 (Sigma Aldrich), hypoxanthine (TCI America), adenine, thymidine and glycine (Sigma Aldrich), sodium formate (Thermo Scientific), ethyl ester GSH, and pralatrexate (Cayman Chemical).

### Incucyte Proliferation Assays

Two thousand five hundred cells per well were plated in a 96-well plate (Corning) in 75 µL of RPMI1640/ 10% dialyzed FBS and doxycycline, and maintained under standard conditions. Plates were inserted into an Incucyte^®^ Live-Cell Analysis System and confluence was measured utilizing live-cell time-lapse imaging.

### Viability assays

Cells were plated in clear bottom, black wall 96-well plates in medium with dialyzed FBS (Biowest). Treatments were added 24 hours after plating. Alamar Blue was directly added to the culture medium of treated and control cells after 72 hours of treatment. Fluorescence was measured using a plate reader (excitation 530 nm, emission 590 nm). Statistical analysis and calculation of EC50 values were performed using GraphPad Prism (RRID:SCR_002798).

### Cell proliferation assays

Cells were plated in 12-(5,000 cells) or 6-well plates (10,000 cells) in RPMI1640 medium with dialyzed FBS. Treatments were added 24 hours after plating. At the end of the experiment, cells were trypsinized and counted using a Coulter particle counter (Beckman).

### Colony formation assay

1,000 cells were plated in each well of a 12-well plate. Treatments were added 24 hours after plating. Medium was replaced every 72 hours. Colonies were fixed with 10% neutral buffered formalin and stained with 0.01% crystal violet (Sigma-Aldrich). For quantitation, crystal violet was solubilized in 30% acetic acid and absorbance at 560 nm was determined using a plate reader.

### LDH release assay

Cells were treated as described above (viability assay). After 72 h, lactate dehydrogenase (LDH) release was assessed using a commercial kit, according to the manufacturer’s instructions (LDH-Cytox Assay Kit, Biolegend).

### Cell Cycle analysis

For cell cycle analysis, cells were treated with DMSO or inhibitors (SHIN1, 5 µM; AGF347, 2 µM; Pralatrexate, 1 nM), harvested after 48 h by trypsin treatment and fixed in 75% ethanol in ice for 4h h. After washing with PBS, cells were stained with FxCycle PI/RNAse staining solution (Invitrogen), and DNA content was measured using an Aurora system (Cyteck).

### 3D spheroid assay

200 cells were plated in 96-well round bottom ultra-low attachment plates (Corning) in RPMI1640 with dialyzed FBS and centrifuged at 350 rpm for 5 minutes. Treatments were added 3 days after plating. Spheroids were microphotographed before treatment and 96h after treatment, and volume was measured using ImageJ (RRID:SCR_003070).

### Western blot

Cells were homogenized on ice in radioimmunoprecipitation assay (RIPA) buffer supplemented with Halt Protease and Phosphatase Inhibitor cocktail (ThermoFisher Scientific). Protein concentrations were determined using the Pierce BCA protein Kit (ThermoFisher Scientific). Western blot analysis was conducted using 50 µg of proteins on ExpressPlus precast gels (Genescript). Proteins were blotted onto polyvinylidene difluoride membranes (Millipore). The membranes were probed with the following antibodies): SHMT1 (Cell Signaling Cat# 80715, RRID:AB_2799957), SHMT2 (Cell Signaling Cat# 33443, RRID:AB_3683566, and actin (Santa Cruz Biotechnology Cat# sc-47778, RRID:AB_62663). All the primary antibodies were used at the vendor-suggested dilution in 5% bovine serum albumin (BSA) in TBS-T. Signals were detected with HRP-conjugated secondary antibodies (ThermoFisher Scientific) and the chemiluminescence substrate Luminata Crescendo (EMD Millipore).

### Metabolomics

SK-ES1 cells (1 million cells/60-mm dish) were seeded in 4 60-mm dishes in 5 ml of complete RPMI 1640 [10% dialyzed FBS (dFBS)]. The cells were allowed to adhere for 24 h. The cells were treated with 10 µM SHIN1, 10 nM pralatrexate (PRTX), or a comparable volume of vehicle (1 % DMSO), in the presence of adenosine (60 μM) and thymidine (10 μM). After 16 h, the cells were washed with PBS (3x); the media was replaced with complete folate- and serine-free RPMI 1640 (containing 130 μM glycine, 2.3 μM folic acid) supplemented with 10% dFBS and [2,3,3-^2^H]serine (250 µM), including drug or vehicle, as appropriate, in the presence of adenosine (60 μM) and thymidine (10 μM). The cells were incubated for 24 h. The media was aspirated, and cells were washed (3x) rapidly (< 30 s) with 5 mL ice-cold PBS; metabolism was quickly quenched with cold 800 µl methanol:water (80:20). The dishes were placed on a shaker and allowed to rock on dry ice for 10 min to cover the entire dish with cold 80:20 methanol:water to extract the metabolites, then harvested by scraping and pipetting the contents into 1.5 mL Eppendorf tubes. The tubes were centrifuged (4°C, 14,000 RPM, 10 min). The supernatants were collected and analyzed by reverse-phase ion-pairing chromatography coupled with negative-mode electrospray-ionization high-resolution mass spectrometry on a stand-alone Orbitrap (ThermoFisher Exactive). Raw metabolite values were adjusted to correct for normal ion distributions and normalized to total proteins from the post-extraction pellet by solubilizing with 500 µl 0.5N NaOH and using 20 µl for protein assay with the Folin-phenol method for protein quantification.

### In vivo studies

6-8 week-old NOD-*scid IL2r*^*null*^ (NSG) mice (JAX #005557; RRID:IMSR_JAX:005557) mice were injected subcutaneously with 3×10^6^ SK-ES-vector or SK-ES-ish*SHMT2* cells. Doxycycline hyclate was dissolved in drinking water (2 mg/ml) and replaced every three days. Tumor volume was calculated from two-dimensional measurements using the equation: tumor volume = (length × width^2^) × 0.5, every three days. Tumor weights were measured at the end of the experiment. Data were plotted and analyzed using GraphPad Prism. All the animal studies were approved by the Einstein Institutional Animal Care and Use Committee.

### Statistical analyses

All data reflect at least three biological replicates, each with three to six technical replicates. Statistical analyses were done in GraphPad Prism using unpaired Student t test or 2-way ANOVA as appropriate.

### Data availability

The data generated in this study are available upon request from the corresponding authors.

## Results

### High expression of one-carbon metabolism genes is associated with poor survival in EWS

Analysis of an EWS gene expression dataset (GSE17679) (13) annotated with clinical and survival data revealed that patients with tumors expressing high levels of RNAs for *SHMT2* or *MTHFD2*, which encode two key enzymes involved in the mitochondrial arm of the one-carbon metabolic pathway (Fig. 1A), experience dramatically reduced overall survival, while no such association was found for *SHMT1* and *MTHFD1*, encoding the corresponding cytoplasmic enzymes, or *MTHFD1L*, which catalyzes the third step in mitochondrial one-carbon metabolism from serine to formate (Fig. 1A, B, C and Suppl. Fig. 1). Notably, both *SHMT2* and *MTHFD2* are expressed at significantly higher levels in recurrent lesions compared to primary tumors (Fig. 1D). These data strongly suggest that increased expression levels and, consequently, activity of enzymes in the mitochondrial arm of the one-carbon metabolism are associated with EWS tumor growth and aggressiveness.

### Genetic downregulation of SHMT2 impairs proliferation of EWS cells

To define the biological consequences of the overexpression of one-carbon metabolism genes, we generated two EWS cell lines (SK-ES1, TC71) expressing control shRNAs or shRNAs targeting *SHMT1* or *SHMT2*. Notably, while we could readily generate cell lines constitutively expressing sh*SHMT1*, EWS cells did not survive constitutive *SHMT2* targeting, such that we had to use a doxycycline-inducible vector for expressing these shRNAs (Fig. 2A). shRNA-mediated depletion of SHMT1 did not alter the proliferation of EWS cells. Knockdown of SHMT2 expression, however, significantly impaired cell proliferation (Fig. 2B and Suppl. Fig. 1).

**Figure 2.**
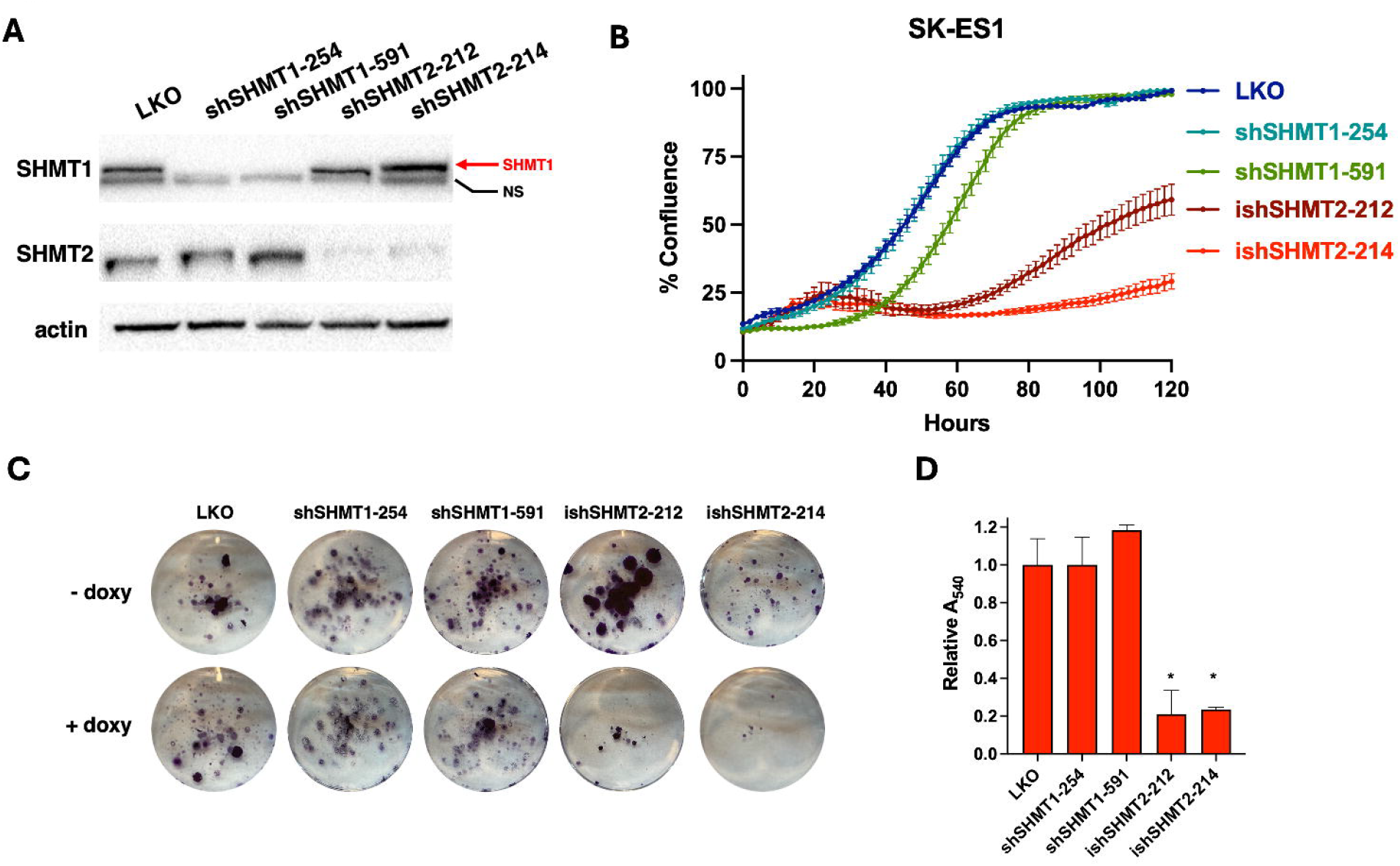
Depletion of *SHMT2*, but not *SHMT1*, impairs proliferation of EWS cells. A) shRNA-mediated depletion of *SHMT1* (constitutive, shSHMT1) or *SHMT2* (inducible, ishSHMT2) in SK-ES1 cells. The arrow points to the specific SHMT1 band, while a non-specific band is indicated by NS. B) Incucyte live analysis of cell proliferation upon depletion of *SHMT1* and *SHMT2* with two shRNA each. LKO: empty vector. C) Representative colony forming assays of SK-ES1 cells upon depletion of SHMT1 or SHMT2. D) Quantitative results from the experiment in C. Asterisks indicate P<0.01.

To assess the long-term impact of genetic inhibition of the one-carbon pathway, we performed colony-forming assays using the SK-ES1 cell line. In line with the results of the short-term cell proliferation assays, shRNA-mediated depletion of SHMT1 did not alter the ability of EWS cells to form colonies. Conversely, depletion of SHMT2 dramatically inhibited colony formation (∼80%, Fig. 2C). The inability to survive in the prolonged absence of SHMT2 suggests that EWS cells are “addicted” to elevated activity of the mitochondrial one-carbon metabolic pathway from SHMT2.

### Pharmacological inhibition of the one-carbon metabolism impairs proliferation of EWS cells

In addition to the classical antifolates, methotrexate and pemetrexed, which primarily target DHFR and thymidylate synthase, respectively (14,15), a number of novel selective inhibitors of enzymes involved in one-carbon metabolism have been developed in recent years (16). Of particular relevance to this study, SHIN1 was synthesized as an inhibitor of SHMT1/2 (17), and AGF347 was developed as a multi-targeted one-carbon inhibitor at SHMT2 in mitochondria with additional inhibition of SHMT1 and de novo purine biosynthesis at glycinamide ribonucleotide (GAR) formyltransferase and 5-aminoimidazole-4-carboxamide ribonucleotide (AICAR) formyltransferase in the cytosol (18,19). SHIN1, a very specific tool compound, has been shown to reduce cell proliferation in various tumor types, including lymphoma (17), rhabdomyosarcoma (20), bladder (21) and lung cancer (22,23) cell lines. AGF347 has shown promising therapeutic activity in vitro and in vivo, in both pancreatic and ovarian cancer models (18,19). Since SHIN1 and AGF347 are not clinic ready, we also used Pralatrexate, the newest clinically approved antifolate inhibitor of DHFR (24), which targets one-carbon metabolism at a different node. We tested the effects of these three inhibitors on a panel of six EWS cell lines encompassing the spectrum of driver mutations associated with EWS development (Table 1). Both SHIN1 and AGF347 were effective in inhibiting the proliferation of EWS cells, with EC50s in the low micromolar range (Fig. 3A, B). Interestingly, all cell lines responded very similarly to the inhibitors, independent of the mutations accompanying the fusion oncogene EWS-FLI1. Pralatrexate, the most potent antifolate tested, completely suppressed cell growth at low nanomolar concentrations (Fig. 3C and Table 2).

**Table 2.** EC_50_ of the three inhibitors in EWS cell lines.

| **EC<sub>50</sub>** | **SHIN1 (μM)** | **AGF347 (μM)** | **PRTX (nM)** |
| --- | --- | --- | --- |
| **CHLA10** | 3.50±0.07 | 0.23±0.04 | 0.92±0.05 |
| **MHH-ES1** | 7.71±0.05 | 3.22±0.05 | 0.93±0.04 |
| **SK-ES1** | 5.47±0.07 | 1.46±0.04 | 0.97±0.06 |
| **TC32** | 1.14±0.05 | 2.00±0.06 | 2.75±0.06 |
| **TC71** | 4.59±0.04 | 1.53±0.04 | 0.94±0.04 |
| **RD-ES** | 5.23±0.04 | 3.41±0.04 | 0.74±0.07 |

**Figure 3.**
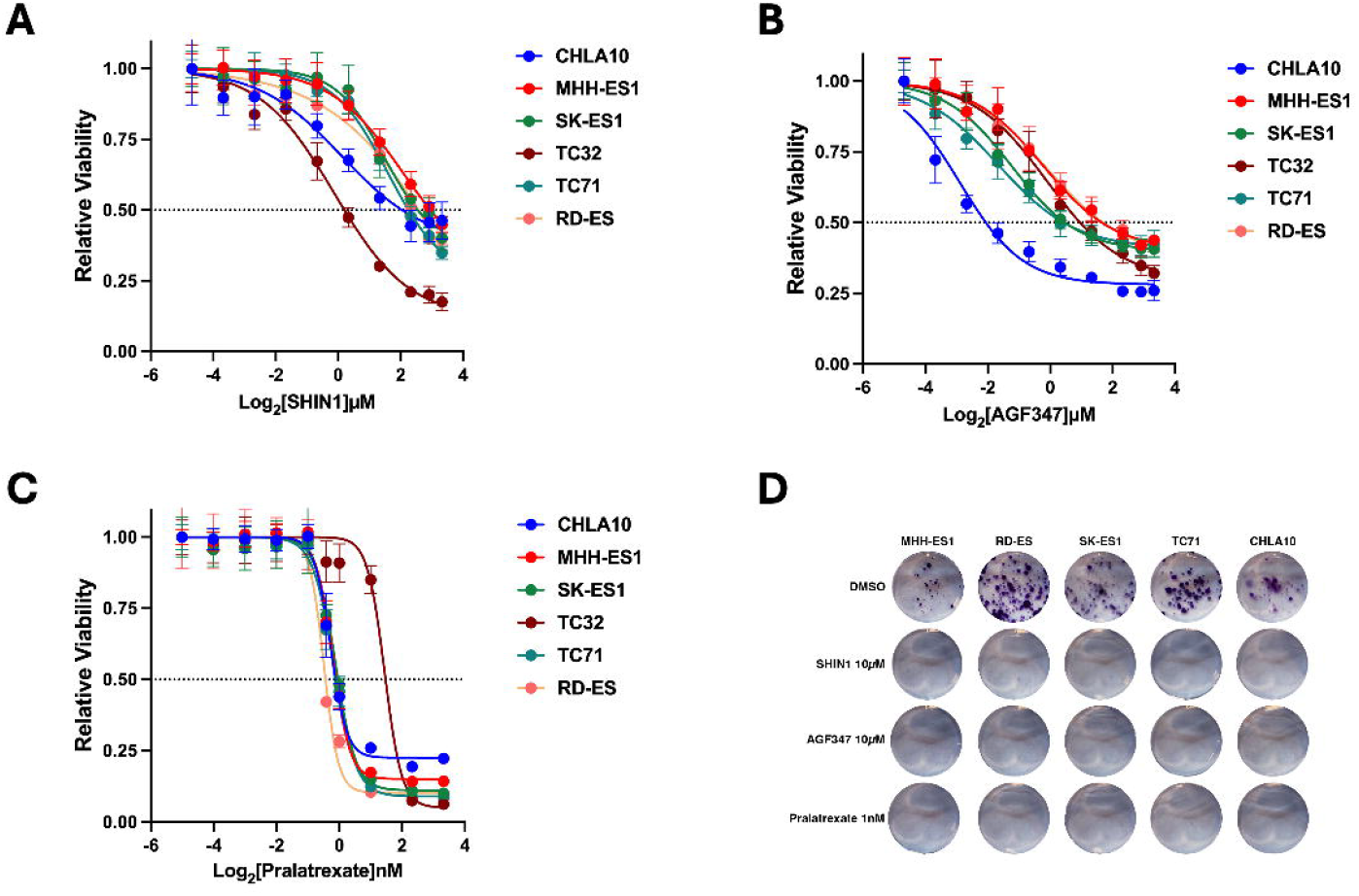
Pharmacological inhibition of one-carbon metabolism impairs EWS cell proliferation. A-C) Viability, measured using alamar blue, of six EWS cell lines treated for 72h with increasing concentrations of SHIN1 (A), AGF347 (B) and Pralatrexate (C). D) Colony forming assay using five EWS cell lines treated for two weeks with the indicated inhibitors.

To assess the effect of long-term pharmacological inhibition of one-carbon metabolism on EWS cell proliferation, we performed colony-forming assays on a panel of EWS cell lines treated with SHIN1, AGF347 or Pralatrexate at concentrations roughly corresponding to their 72h EC_70_. After two weeks, colony formation was completely suppressed by these antifolates (Fig. 3D). These data significantly extend the genetic approaches described in Fig. 2 and further suggest that, in EWS cells, one-carbon pathway inhibition leads to a profound growth suppression.

Next, we tested the effect of one-carbon metabolism inhibition in SK-ES1 3D spheroid models, which recapitulate the biology of solid tumors more closely than classical cell culture in a dish. After three days of drug treatment, we found that both SHIN1 and Pralatrexate inhibited spheroid growth by more than 50%, further validating the efficacy of targeting different nodes of one-carbon metabolism in EWS (Fig. 4A).

**Figure 4.**
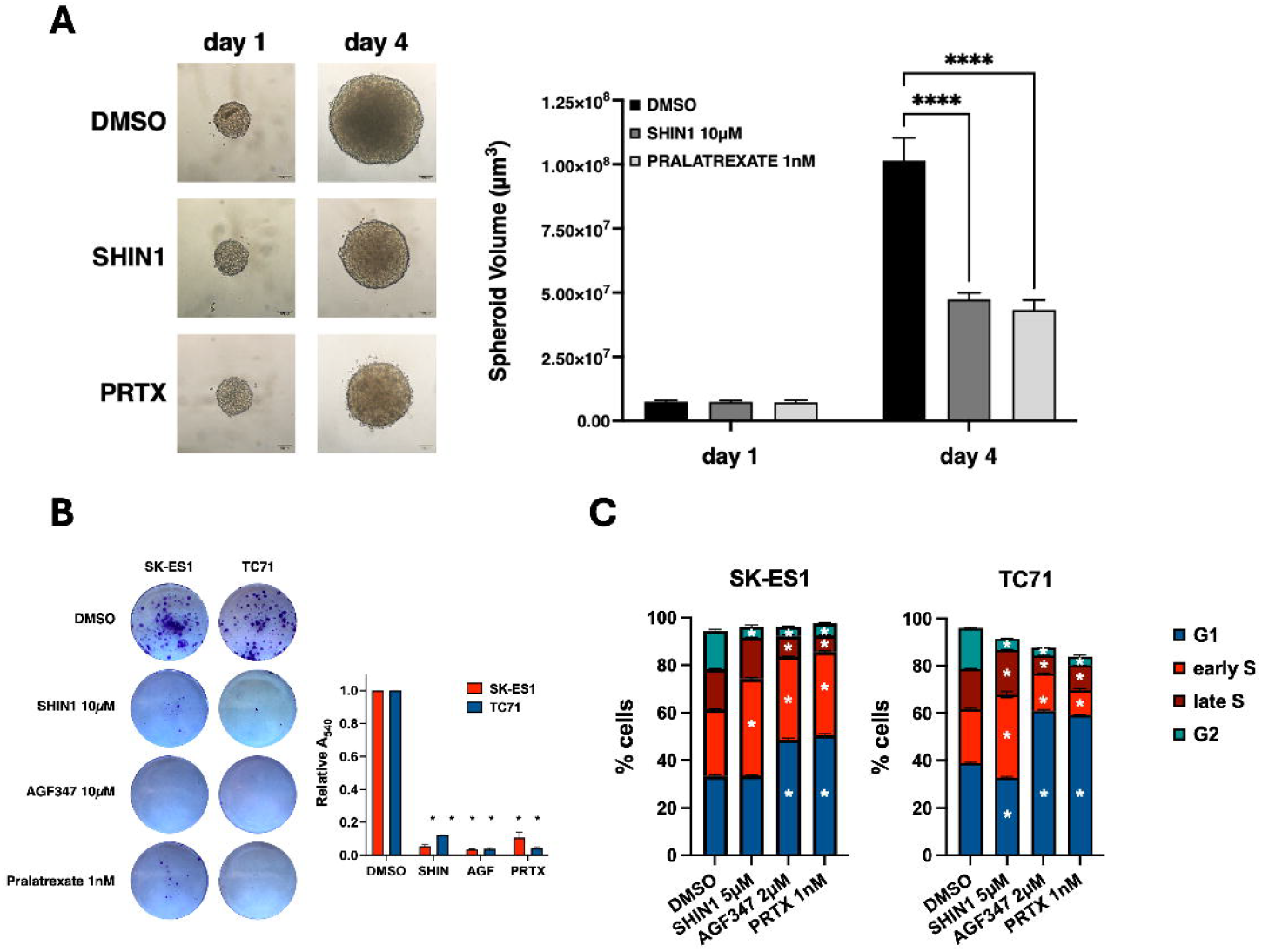
Pharmacological inhibition of one-carbon metabolism results in inhibition of EWS spheroid growth and colony formation resulting in G1-early S phase arrest. A) Analysis of the inhibitory effect of SHIN1 and Pralatrexate on the growth of SK-ES1 spheroids. Asterisks indicate P<0.001. B) Colony forming assay of SK-ES1 and TC71 cells treated with the indicated compounds for six days and then grown without drugs for ten additional days. Quantitative results are shown in the graph to the right. Asterisks indicate P<0.0001. C) Cell cycle analysis of SK-ES1 and TC71 cells treated with the indicated compounds for 48h. Asterisks indicate P<0.01 from replicate experiments.

To determine whether the growth suppression consequent to one-carbon pathway inhibition is reversible upon drug “washout”, we treated SK-ES1 and TC71 EWS cells with one-carbon metabolism inhibitors for six days and then removed the compounds for an additional 10 days. Strikingly, both cell lines almost completely failed to resume proliferation (>80% reduction, Fig. 4B). These results strongly suggest that, in EWS cells, one-carbon pathway inhibition leads to a prolonged state of growth suppression.

To identify the mechanisms leading to growth suppression upon inhibition of the one-carbon metabolic pathway, we measured the effects of SHIN1, AGF347 or Pralatrexate treatment on the induction of cell death using an LDH release assay. We did not find any evidence of cell death upon treatment with these inhibitors (data not shown). We then analyzed the effects of the three inhibitors on the cell cycle distribution of SK-ES1 and TC71 cells and observed a significant reduction of cells in the G2 phase and an accumulation of cells in the G1 and/or early S phases of the cell cycle. These data suggest that inhibition of one-carbon metabolism impairs the ability of EWS cells to complete DNA replication (Fig. 4C).

### One-carbon pathway inhibition leads to nucleotide depletion

Mitochondrial one-carbon metabolism from serine results in the synthesis of glycine (via SHMT2), NADH (via MTHFD2) and formate (via MTHFD1L2) (7-9). Formate passes to the cytosol where it participates in cellular anabolism. Inhibition of the one-carbon pathway at the SHMT2 node is predicted to lead to two critical metabolic deficiencies. These include depletion of formate, which is the principal source of one-carbon units for de novo nucleotide synthesis in the cytosol, and glycine, a critical amino acid for the synthesis of proteins and purine nucleotides (Fig. 1A).

To further define the relative impact of inhibiting one-carbon metabolism on these cytosolic pathways in EWS cells, we first tested whether providing exogenous metabolites can rescue cells from the growth-suppression caused by pharmacological inhibition of SHMT1 and -2. Supplementation of the growth medium with 1 mM formate, in the presence of glycine (133 µM), partially restored the proliferative ability of EWS cells treated with SHIN1 (∼70%) (Fig. 5A) and or AGF347 (∼60%) (Fig. 5B). The additional impact of AGF347 in these experiments likely reflects the direct targeting of de novo purine biosynthesis at GAR and AICAR formyltransferases (18), which are downstream of formate.

**Figure 5.**
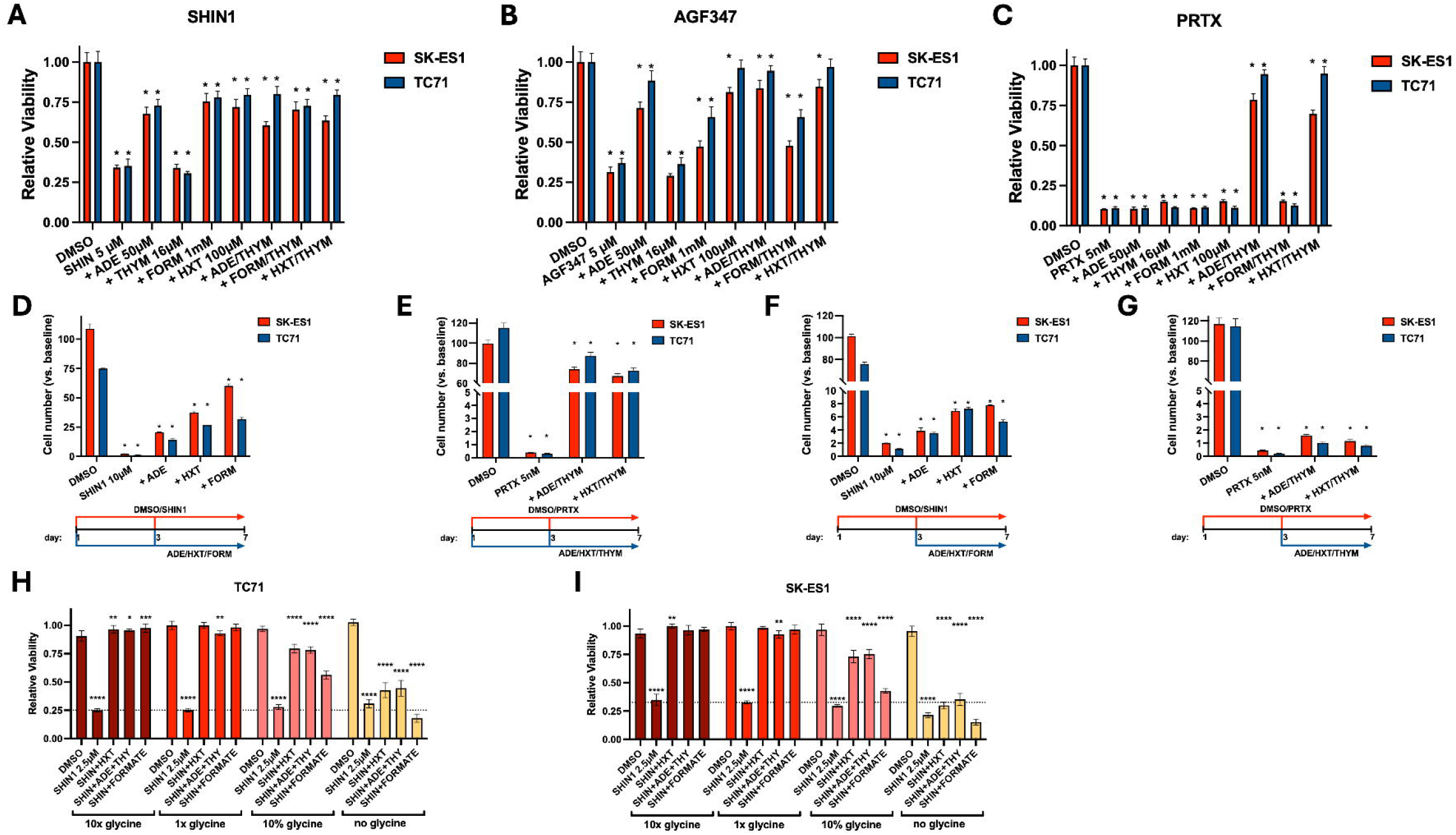
Progressive loss of the ability of relevant metabolites to rescue the proliferation of EWS cells after one-carbon metabolism inhibition. A-C) Rescue by exogenous adenine, thymidine, formate and hypoxanthine addition of the growth inhibition induced by 72 h treatment of SK-ES1 and TC71 cells with one-carbon inhibitors. D, E) Rescue by metabolite supplementation of the growth inhibition induced by seven day-treatment of SK-ES1 and TC71 cells with SHIN1 (D) and Pralatrexate (E). F, G) Rescue effect of delayed (at day 3) metabolite addition on the growth inhibition induced by seven day-treatment of SK-ES1 and TC71 cells with SHIN1 (F) and Pralatrexate (G). A-G) Asterisks indicate P<0.001 vs. DMSO. H, I) Exogenous glycine deprivation impairs the ability of hypoxanthine, adenine, thymidine and formate (concentrations as in A-C) to rescue the proliferation impairment consequent SHMT1/2 inhibition with SHIN1. Glycine-free RPMI1640 was supplemented with 1.33 mM (10x), 133 µM (1x) or 13.3 µM (10%) glycine. *, p<0.05; **, p<0.01; ***, p<0.001; ****, p<0.0001.

To distinguish between the drug effects on one-carbon metabolism leading to purine nucleotide versus thymidylate synthesis, we also tested the protective impact of exogenous adenine or hypoxanthine, which provide substrates for the purine salvage pathway; results were compared to those for exogenous thymidine which is metabolized to thymidylate, thus circumventing the thymidylate synthase reaction. Both adenine and hypoxanthine substantially reversed the proliferative block induced by SHIN1 and AGF347 in EWS cells. This suggests that, in the presence of glycine, purine depletion is a major consequence of inhibiting one-carbon metabolism by these agents (Fig. 5A, B). Thymidine supplementation did not rescue the proliferation deficit and did not alter the extent of the rescue induced by hypoxanthine and adenine (Fig. 5A, B). Thus, thymidylate levels do not appear to be limiting for proliferation of EWS cells treated with SHIN1 or AGF347. In contrast to our findings with SHIN1 or AGF347, rescue from the inhibitory effects of Pralatrexate required a combination of adenine (or hypoxanthine) with thymidine, results entirely consistent with its inhibition of DHFR (Fig. 5C).

To test whether exogenous metabolite supplementation can also prevent the antiproliferative effects of one-carbon metabolism inhibition at SHMT2 or DHFR over a longer period of time, we treated SK-ES1 and TC71 cells with a high dose of SHIN1 (10 µM) or Pralatrexate (5 nM), alone or in the presence of rescuing metabolites, for seven days, with a complete change of medium and compounds after three days. Exogenous adenine, hypoxanthine and formate were only able to partially prevent growth suppression. This strongly suggests that, upon prolonged inhibition of one-carbon metabolism at SHMT1/2, purine depletion is accompanied by deficiencies in additional metabolites or biological processes, which become limiting for EWS cell proliferation (Fig. 5D). The effect of prolonged treatment with Pralatrexate could, in contrast, be prevented to a greater extent by supplementation with adenine (or hypoxanthine) with thymidine (Fig. 5E).

Next, we tested whether the growth suppression caused by inhibition of one-carbon metabolism is reversible after being fully established. To this end, we supplemented purine-generating metabolites three days after treatment with a high dose of SHIN1 and measured cell proliferation after an additional four days of combined treatment. In the presence of adenine, hypoxanthine or formate, EWS cells resumed a low level of proliferation despite continuous SHIN1 presence, indicating that, within the time frame analyzed, purine availability can only partially reverse the pre-existing growth arrest induced by inhibition of one-carbon metabolism at the SHMT level (Fig. 5F). Further extending the pre-inhibition to six days before hypoxanthine supplementation resulted in even lower proliferation recovery, while higher resumption of proliferation was observed if SHIN1 was removed from the growth medium (Suppl. Fig. 3). Inhibition of proliferation with Pralatrexate led to a more profound effect, which was only nominally reversed by metabolite addition after three days of pre-treatment (Figure 5G). These data suggest that inhibition of one-carbon metabolism in EWS cells results in a progressively irreversible growth suppression.

In a previous study, we found that thyroid cancer cells become fully auxotrophic for glycine upon inhibition of SHMT1/2 with SHIN1. In the absence of exogenous (i.e., culture medium) glycine (133 µM), the growth suppressive effect of SHIN1 was dramatically amplified, leading to cell death (25). We tested the effect of removing extracellular glycine on the response of EWS cells to SHIN1. Surprisingly, despite the complete inhibition of glycine synthesis from serine, the growth inhibitory effects of SHIN1 did not change with the presence or absence of glycine in the culture medium, and SHIN1-treated EWS cells survived in glycine-free medium (Suppl. Fig. 4). Additionally, supplementation of the culture medium with 10-fold excess glycine (1.33 mM) did not change the effect of SHIN1 (Fig. 5 H, I). These data suggest that, contrary to results with thyroid cancer cells (25), the growth suppressive effects of one-carbon metabolism inhibitors on EWS cells are largely independent of glycine availability. EWS cells might procure sufficient glycine for basic survival through alternative pathways. However, in the absence of exogenous glycine, supplementation with purine-producing metabolites was unable to restore EWS cell proliferation in the presence of SHIN1, indicating that the hypothetical alternative source of glycine may not be sufficient to meet the demand of all the metabolic processes requiring glycine (Fig. 5 H, I).

### Metabolomic analysis of one-carbon flux

We performed metabolomic analysis of SK-ES1 cells with [2,3,3-^2^H]L-serine to study the impact of drug treatments on intracellular one-carbon metabolism. SK-ES1 cells were treated with vehicle (1 % DMSO), Cells were treated with pralatrexate (10 nM) or SHIN1 (10 µM) for 24 h, followed by [2,3,3-^2^H]L-serine (250 µM) for another 24 h, then processed for determinations of total serine and glycine, along with their isotopomer distributions (M+3, M+2, M+1, and M+0 for serine; M+1 and M+0 for glycine; M+n represents species with n deuterium atoms). The results for the drug-treated cells were compared to those for vehicle-treated cells. A schematic of [2,3,3-^2^H]L-serine metabolism via SHMT2 and SHMT1 (including distributions of deuterium atoms in downstream metabolites including dTTP) is included in Fig. 6A.

**Figure 6.**
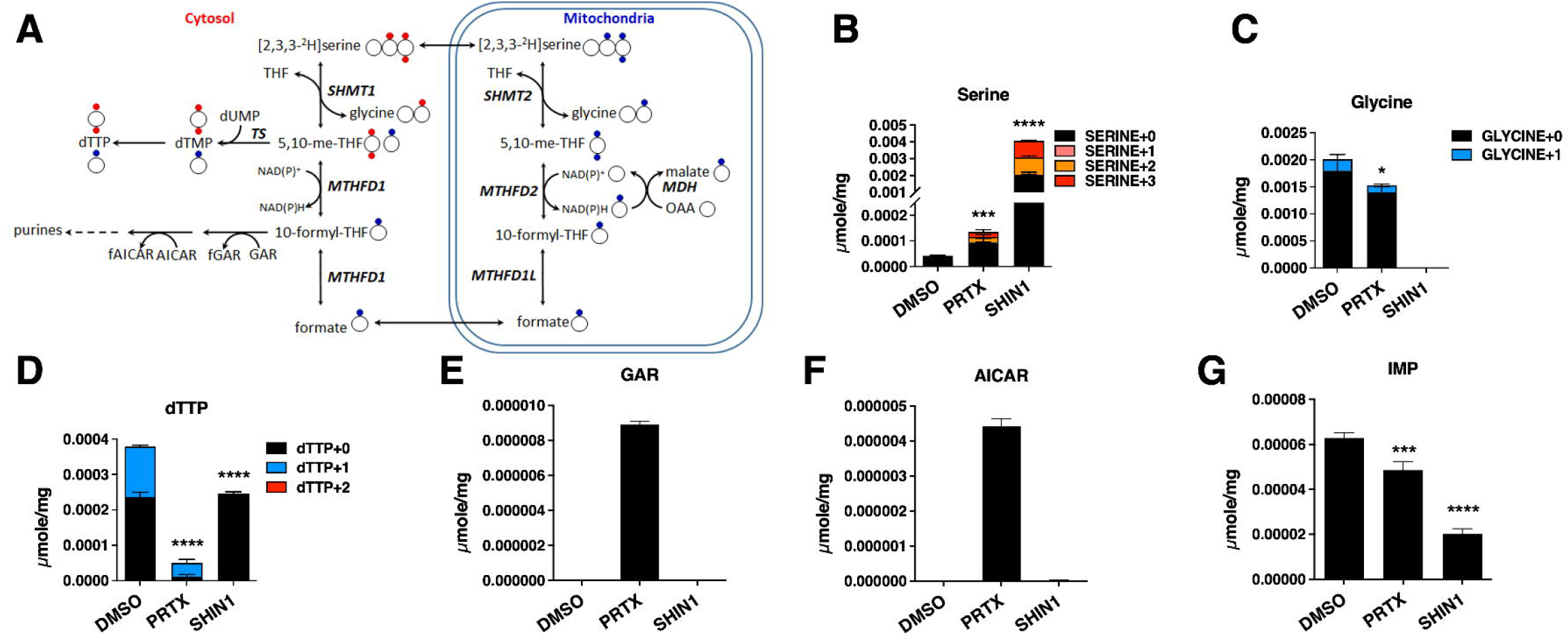
Effects of SHIN1 and Pralatrexate on one-carbon flux through the mitochondrial and cytoplasmic compartments. A) Schematic of one-carbon flux through the mitochondrial and cytosol compartments using [2,3,3-^2^H]L-serine. The ^2^H atoms in [2,3,3-^2^H]L-serine (blue circle) in the mitochondria are metabolized to generate [^2^H]formate. [^2^H]Formate in the cytosol is converted to [^2^H]10-formyl THF and [^2^H]5,10-methylene THF, resulting in [^2^H]dTMP and [^2^H]dTTP (as M+1). Reversal of SHMT1 activity in the cytosol results in ^2^H atoms in [2,3,3-^2^H]serine (red) being metabolized to [^2^H]5,10-methylene THF, which is utilized by TS to synthesize [^2^H]dTMP and [^2^H]dTTP (as M+2). B, C) Total serine (B) and glycine (C) and the serine-isotopomer distributions in SK-ES1 cells treated with vehicle (DMSO), 10 nM Pralatrexate (PRTX) or 10 µM SHIN1. The increase in the M+1 and M+2 serine isotopomers in the presence of the inhibitors is a direct reflection of accumulated [2,3,3-^2^H]L-serine accompanying the loss of SHMT2 activity, combined with low levels of one-carbon flux through SHMT2 and MTHFD2 in mitochondria and SHMT1 in the cytosol. D) total dTTP pools are shown with the isotopomer distributions (M+0, M+1, M+2) for untreated and inhibitor-treated SK-ES1 cells compared to DMSO-treated cells. E-G) GAR (E), AICAR (F) and IMP (G) pools were measured in vehicle and inhibitor-treated cells. Unpaired t test was performed for total serine, total glycine, total dTTP and IMP, in comparison to DMSO-treated sample. *, p<0.05; ***, p<0.001; ****, p<0.0001.

Treatment of SK-ES1 cells with Pralatrexate and SHIN1 increased total serine levels 3-fold and 80-fold, respectively, above the levels in vehicle-treated cells (Fig. 6B). For SHIN1 treated cells, elevated serine pools were accompanied by increased M+3 (180-fold) and M+2 (449-fold) serine isotopomers over vehicle-treated cells. There was ∼24 % decreased total glycine for cells treated with Pralatrexate. Strikingly, SHIN1 treatment resulted in a complete loss of glycine (Fig. 6C).

The ^2^H-dTTP isotopomer distribution from [2,3,3-^2^H]L-serine was used as a direct measure of relative mitochondrial and cytosolic one-carbon fluxes through SHMT2 and SHMT1 into dTMP (and dTTP) (10). Thus, the ^2^H flux from [2,3,3-^2^H]L-serine to [^2^H]formate via SHMT2 in mitochondria results in M+1 [^2^H]dTTP, whereas the ^2^H flux from [2,3,3-^2^H]L-serine via reversal (serine-to-glycine) of SHMT1 catalysis in cytosol results in M+2 [^2^H]dTTP (Figure 6A) (7,10,17,18). As expected, treatment of SK-ES1 cells with SHIN1 decreased total dTTP pools (35 %) and M+1 dTTP was completely abolished compared to vehicle-treated cells (Fig. 6D). These results are entirely consistent with direct targeting of SHMT2 by SHIN1 (17). Meanwhile, Pralatrexate treatment resulted in a dramatic decrease of both the dTTP+0 (95 %) and dTTP+1 (73 %), most likely due to the potent inhibition of DHFR resulting in near complete loss of THF with downstream effects on one-carbon transfer. Interestingly, there was no detectable M+2 dTTP, suggesting that the SK-ES1 cells rely primarily on mitochondrial one-carbon metabolism for proliferation.

We also assessed the impact of Pralatrexate and SHIN1 on de novo purine biosynthesis in SK-ES1 cells. We measured accumulated GAR and AICAR (respective substrates for GARFTase and AICARFTase, respectively; Figure 6A) by LC-MS. GAR and AICAR levels were nominal in vehicle- and SHIN1-treated cells but were accumulated to significant levels in Pralatrexate-treated cells (Fig. 6E, F). Although no accumulation of GAR and AICAR was observed for either vehicle- and SHIN1-treated SK-ES1 cells, the basis for this result is likely very different, with normal purine biosynthesis ongoing for the former and a complete shutdown of purine pathway resulting from glycine depletion for the latter. This is reflected in the levels of IMP (Fig. 6I). Inhibition of DHFR by Pralatrexate profoundly decreased purine biosynthesis, reflected in substantial accumulation of GAR and AICAR intermediates. Pralatrexate and SHIN1 treatments decreased total IMP pools by 23 % and 68 %, respectively (Fig. 6G).

Thus, inhibition of one-carbon metabolism in EWS cells leads to a near-complete cessation in the synthesis of glycine and purines with SHMT1/2 inhibitors, and a significant impairment with DHFR inhibition, which has its major impact on dTTP synthesis.

### Effect of SHMT2 depletion on tumor growth in vivo

To assess in vivo the effect of inhibiting mitochondrial one-carbon metabolism in EWS cells, we used SK-ES1 xenografts expressing either a control shRNA or shSHMT2 (doxycycline-inducible) into NSG mice. Mice were injected subcutaneously on one flank with the control cells and on the other flank with the SHMT2-depleted cells. The SHMT-depleted cell lines showed a dramatic reduction in tumor growth rate (Fig. 7A) as well as in tumor weight at the end of the experiment (4.74 g for vector-transduced cells vs. 0.38 g for ish*SHMT2*-expressing cells, Fig. 7B), thus providing direct evidence that inhibition of one-carbon metabolism could be a viable treatment strategy for EWS patients.

**Figure 7.**
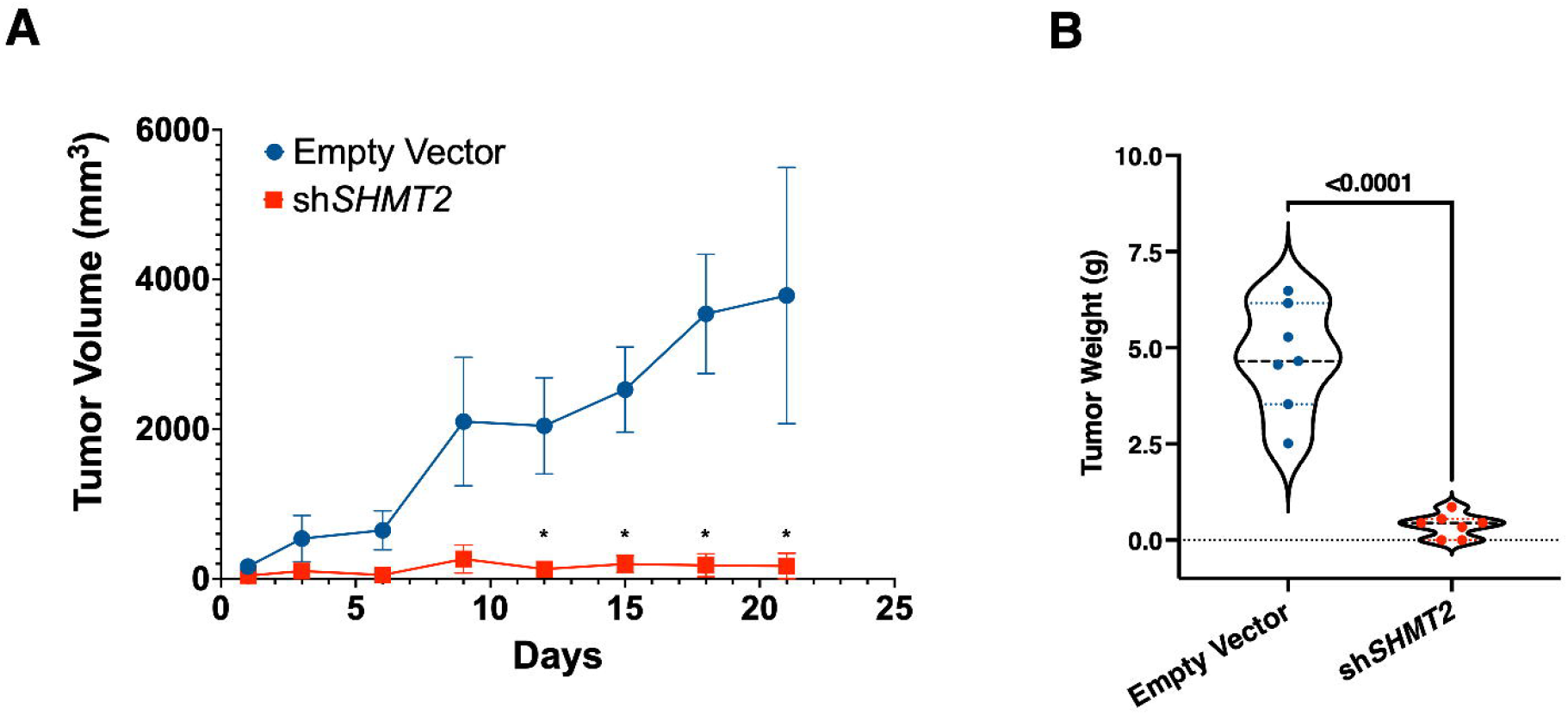
In vivo efficacy of genetic one-carbon metabolism inhibition. A) shRNA-mediated depletion of *SHMT2* severely impairs in vivo tumor growth in SK-ES1 xenografts, resulting in significantly smaller tumors, as shown by the difference in weight of the excised tumors at the end of the experiment (B). Asterisks in A indicate P<0.01.

## Discussion

Cancer cells undergo various metabolic alterations to adapt to their heightened proliferative demands and their unique microenvironment. These changes enable tumors to sustain rapid growth, survive under conditions of limited oxygen or nutrient supply, and resist stress from their surroundings. These metabolic adaptations frequently create vulnerabilities that are unique to cancer cells, distinguishing tumors from normal cells. As such, they represent valuable targetable liabilities that can be exploited in the development of novel therapeutic strategies (26).

One of the metabolic pathways most often upregulated by tumor cells is one-carbon metabolism, which utilizes serine and dietary folates to generate glycine and tetrahydrofolate (THF)-bound one-carbon units, required for de novo nucleotide biosynthesis, NAD(P)H and glutathione production, methionine regeneration, DNA methylation and translation of mitochondrial proteins (7-9).

*SHMT2* and *MTHFD2*, encoding two key enzymes of the mitochondrial arm of the one-carbon pathway, have been reported to be among the most consistently upregulated metabolic genes in cancer (11,12). Not surprisingly, this has led to renewed interest in targeting mitochondrial one-carbon metabolism for cancer (17,22,27,28).

Selective inhibitors of these enzymes, however, are still at an early stage of preclinical development, while the value of repurposed inhibitors with reported activity against SHMT2 (e.g., sertraline (29), metformin (30)) is difficult to assess due to lack of target specificity. Pralatrexate, a clinically approved antifolate drug targeting DHFR (24), offers however the possibility to test the efficacy of targeting one-carbon metabolism, although at a different node.

In this study, our data shed important new light on the role of one-carbon metabolism in driving the proliferation of EWS cells and demonstrate a remarkable dependence on this pathway, a notion supported by the dramatic effect of genetic and pharmacological inhibition of one-carbon metabolism. Inhibition of SHMT1/2 or DHFR both led to strong cytostatic effects that persisted for several days even when the inhibitor is removed. Whether these cells undergo irreversible senescence or enter a prolonged but still partially reversible state of quiescence remains to be addressed in future work.

When glycine is present, metabolites such as hypoxanthine and adenine, but not thymidine, rescue the proliferation block associated with inhibition of one-carbon metabolism at the level of SHMT1/2. While in agreement with other studies, where purine synthesis is the specific essential conduit of SHMT1/2 inhibition (17), these data raise the question of why thymidylate does not become limiting, at least in the time frame analyzed, as it does when one-carbon metabolism is inhibited at the level of DHFR with Pralatrexate.

In previous studies, we have shown that inhibition of one-carbon metabolism at SHMT1/2 in anaplastic thyroid cancer cells leads to glycine auxotrophy and rapid cell death when cells are deprived of extracellular glycine (25). Interestingly, this does not seem to be the case for EWS cells, since the presence or absence of glycine in the medium does not alter the efficacy of both SHIN1 and AGF347. While it is possible that alternative intracellular sources of glycine (31) are able to maintain viability of EWS cells when both one-carbon metabolism-mediated synthesis and extracellular uptake are impaired, the inability of these cells to remain proliferative when adenine or hypoxanthine are supplemented suggests that this alternative source of glycine is still limiting, a notion that offers an interesting targetable liability for future therapeutic applications.

We used a genetic approach to inhibit one-carbon metabolism in vivo in an immunocompromised xenograft model. Consistent with our in vitro findings, our in vivo results strongly suggest that SHMT inhibition is a viable and promising strategy to on which to build novel therapeutic approaches to EWS. While inhibition of one-carbon metabolism in EWS cells appears to be cytostatic, the prolonged growth arrest observed in cell culture suggests the possibility of extended tumor control in vivo, a therapeutic goal of utmost importance in advanced, metastatic or recurrent EWS management. In addition, we believe that these approaches represent a solid foundation for the development of rationally designed synthetic lethal strategies harnessing the liabilities exposed by one-carbon metabolism impairment.

## Acknowledgements

Research reported in this publication was supported by NIH grants CA128943 to ADC and CA53535 to LM and ZH and the Eunice and Milton Ring Endowed Chair for Cancer Research (to LM). Additional support was received through grants from the National Pediatric Cancer Foundation and Alan B. Slifka Foundation to ADC and DML, and the V Foundation for Cancer Research to DML. We acknowledge the Animal Housing and Flow Cytometry Core Facilities of Albert Einstein College of Medicine, which are partially supported by the NIH Cancer Center Support Grant to the Albert Einstein Cancer Center (P30CA013330). The Pharmacology and Metabolomics Core at the Barbara Ann Karmanos Cancer Institute was supported in part by the NIH Support Grant P30CA22453.

## Declaration of competing interest

The authors declare that they have no known competing financial interests or personal relationships that could have appeared to influence the work reported in this paper.

## Supplementary Figures

**Suppl. Fig. 1**

Overall survival of EWS patients (n=88) expressing high or low levels of *SHMT1, MTHFD1, MTHFD1L*. Analysis was performed on dataset GSE17679 using R2: Genomics Analysis and Visualization Platform (http://r2.amc.nl).

**Suppl. Fig. 2**

Incucyte live analysis of TC71 cell proliferation upon depletion of *SHMT1* and *SHMT2*.

**Suppl. Fig. 3**

Effect (at day 10) of delayed hypoxanthine addition or SHIN1 removal, both at day 6, on the growth inhibition induced in SK-ES1 and TC71 cells SHIN1 treatment.

**Suppl. Fig. 4**

Exogenous glycine deprivation does not alter the efficacy of 72h treatment of EWS cells with SHIN1. Cells were grown in glycine-free RPMI1640 with or without supplementation with 133µM glycine.

## Notes

### Competing Interest Statement

The authors have declared no competing interest.

### Summary of Updates

Funding information and edited author affiliations

