## Supplementary figures and images for "Inhibition Of One-Carbon Metabolism In Ewing Sarcoma Results In Profound And Prolonged Growth Suppression Associated With Purine Depletion"

### SF1

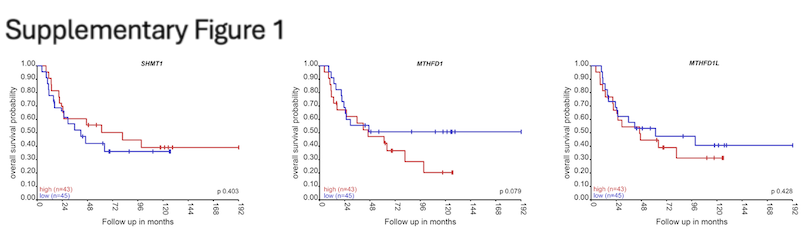

### SF2

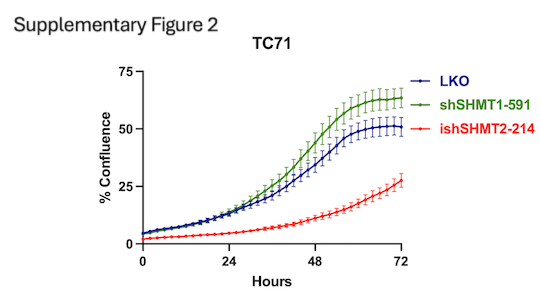

### SF3

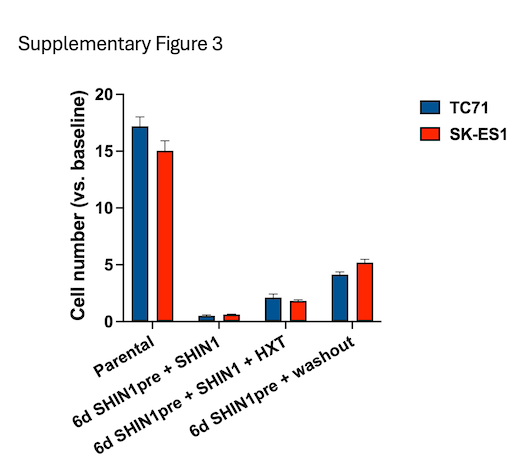

### SF4

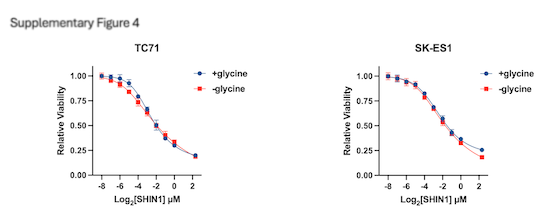
